# Viral protease-Initiated Pyroptosis Activator mRNA therapy as a Universal Antiviral Strategy

**DOI:** 10.1101/2025.11.14.688189

**Authors:** Lin Li, Xiu-Li Yan, Hao-Yang Wang, Hang-Yu Zhou, Yu-Yan Li, Jian Li, Rong-Rong Zhang, Lu Lv, Xing-Yao Huang, Hao Du, Tian-Shu Cao, Qing Ye, Hui Zhao, Yan Fu, Aiping Wu, Yi-Jiao Huang, Cheng-Feng Qin

## Abstract

Although therapeutic drugs targeting gasdermin (GSDM)-mediated pyroptosis have made remarkable progress in treating various diseases, their potential for antiviral therapy remains largely unexplored. Inspired by the autoinhibitory mechanism of GSDM proteins and the clinical success of mRNA vaccines, herein we propose a concept of viral protease-initiated pyroptosis (VIP) mRNA therapy as a universal antiviral strategy. Using Hepatitis A virus (HAV) as a model, we rationally engineered an HAV-specific VIP activator (VIPA) by replacing the native gasdermin D (GSDMD) cleavage motif with that of HAV 3C protease. Expression of this VIPA selectively triggered pyroptosis in HAV-infected cells, thereby terminating viral replication. Remarkably, *in vivo* delivery of VIPA mRNA *via* lipid nanoparticles (LNPs) significantly suppressed viral replication and shedding, alleviated liver injury, and elicited a pyroptosis-driven antiviral immune response in uninfected cells. Similarly, VIPA against Zika virus (ZIKV) also exhibited potent antiviral efficacy in both cell culture and mouse models. Furthermore, a generative artificial intelligence (AI) platform was deployed to design *de novo* cleavage motifs for the SARS-CoV-2 main protease NSP5, together with experimental screening, yielding optimized VIPAs with enhanced antiviral potency. Our work establishes the concept of VIPA mRNA therapy as a versatile platform for the treatment of diverse viral infections *via* "kill-and-alert" mechanism.

## Introduction

Pyroptosis is a form of programmed lytic cell death mediated by the gasdermin (GSDM) protein family (*1*, *2*). Upon recognition of exogenous or endogenous signals, GSDMs become activated to form membrane pores, causing loss of membrane integrity and release of pro-inflammatory cytokines as well as abundant cellular contents, including pathogen-associated molecular patterns (PAMPs) and damage-associated molecular patterns (DAMPs) (*3–5*). GSDM activation and the resulting pyroptosis have been extensively studied across a broad spectrum of physiological and pathological processes, such as infection, cancer, sepsis, asthma, and autoimmune disease (*6–10*). Thus, GSDM-mediated pyroptosis has emerged as a promising therapeutic target, and a number of GSDM inhibitors have been identified or rationally designed with promising therapeutic efficacy in preclinical disease models and clinical trials (*11–13*).

However, the role of GSDM-mediated pyroptosis during viral infection remains largely under investigated. In principle, pyroptosis serve as a host defense mechanism against viral infection, disrupts viral replication niches, and recruits additional immune cells to eradicate the infection (*14*). During human rotavirus infection, host Nlrp9b inflammasome promotes GSDMD-induced pyroptosis to restrict viral spread within the intestinal mucosa (*15*). However, excessive or dysregulated GSDM activation can be detrimental. GSDMD-dependent pyroptosis has been shown to exacerbate hyperinflammation and immunopathology during SARS-CoV-2 and influenza A virus (IAV) infections, thereby aggravating disease severity (*16*, *17*). Zika virus (ZIKV) and IAV infections trigger GSDME mediated pyroptosis, and silencing GSDME expression alleviates virus-induced tissue damage (*18*, *19*). These diverse mechanisms and context-specific roles of pyroptosis in individual viral infections make it extremely challenging to develop a universal strategy to precisely orchestrate pyroptosis of virus-infected cells.

Viruses encode their own viral proteases, which process the polyproteins into individual viral protein within infected cells. These proteases are typically translated early during infection and exhibit high specificity for their respective cleavage motifs (*20*, *21*), making them attractive antiviral targets. In parallel, GSDM proteins are maintained in an autoinhibited conformation in resting cells, and pyroptosis is normally triggered upon proteolytic cleavage of their interdomain linker by host inflammatory caspases (*1*, *2*). To mimic host caspase initiated pyroptosis, we proposed the concept of Viral protease-Initiated Pyroptosis (VIP) to specially kill the infected cells by replacing the native caspase-cleavage motif in GSDMs with that of a specific viral protease (Fig. 1A). Accordingly, upon viral infection, the early-translated viral protease would precisely cleave and activate the engineered GSDMs, leading to rapid pyroptosis in infected cells, and release of viral and cellular antigens as alarming signals.

**Fig. 1.**
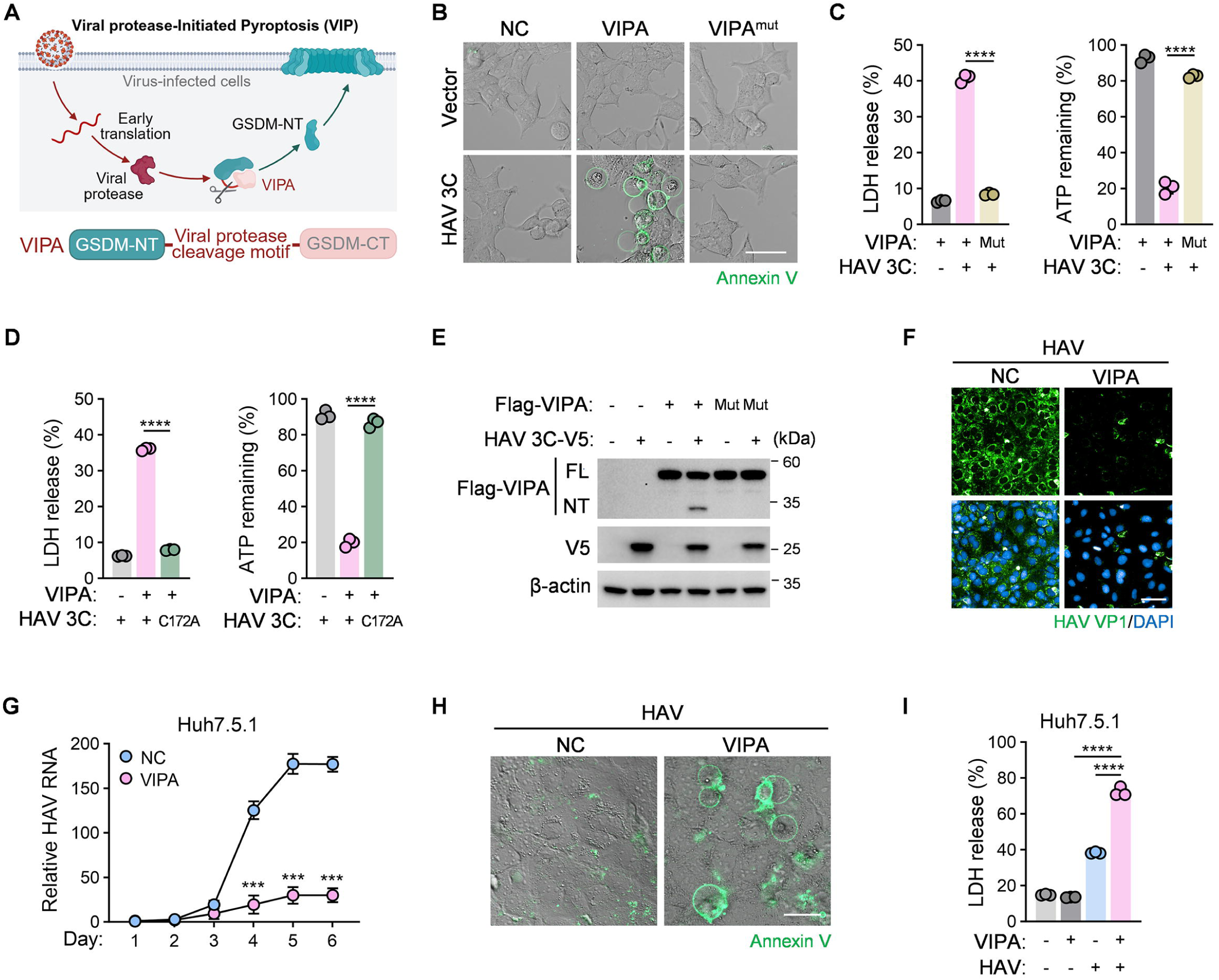
Design and validation of VIPA against HAV. (**A**) Schematic illustration of the VIP concept and VIPA construct, created with BioRender.com. (**B to E**) HEK293T cells (n = 3) were co-transfected with HAV 3C protease or its active-site mutant (C172A), together with VIPA or its protease-resistant mutant (VIPA^mut^). (**B**) Live-cell imaging showing FITC-Annexin V staining (scale bar, 20 μm). (**C and D)** Quantification of LDH release and intracellular ATP levels at 12 h post-transfection. (**E)** Immunoblot analysis of full-length and cleaved VIPA, along with HAV 3C, at 24 h post-transfection. (**F to I)** Huh7.5.1 cells (n = 3) were infected with HAV and treated with or without VIPA. (**F)** Immunofluorescence staining of HAV VP1 (green) and nuclei (DAPI, blue) in Huh7.5.1 cells at 4 dpi following VIPA treatment (scale bar, 50 μm). (**G)** Time course of intracellular HAV RNA levels in Huh7.5.1 cells with or without VIPA treatment from 1 to 6 dpi. (**H)** Live-cell imaging of Huh7.5.1 cells at 4 dpi, infected with HAV and treated with or without VIPA, stained with FITC-Annexin V (scale bar, 20 μm). (**I)** LDH release in HAV-infected Huh7.5.1 cells at 4 dpi, treated with or without VIPA. Data are presented as mean ± standard deviation (SD) of biological replicates.

Herein, to translate the VIP concept into feasible antiviral therapy, we rationally designed and developed VIP activators (VIPAs) mRNA therapy with the established mRNA-LNP platform (*22*), with robust antiviral efficacy in multiple viral infection models. Comprehensive mechanistic investigation combined with generative artificial intelligence (AI) modeling highlight the great potential of this universal "kill-and-alert" antiviral mRNA therapy.

### Rational design of viral protease-initiated pyroptosis activator (VIPA)

To implement the VIP concept, we first designed VIPA by replacing the native caspase-1 cleavage motif (FLTD↓GV) of GSDMD with the recognition motif of HAV 3C protease (LRTQ↓SF) (Fig. 1A). A scrambled construct (VIPA^mut^, QLTSFR) was used as a negative control. In HEK293T cells, co-expression of VIPA and 3C protease triggered hallmark features of pyroptosis, including pronounced cell swelling observed by live-cell imaging (Fig. 1B), lactate dehydrogenase (LDH) release, and ATP depletion (Fig. 1C). These effects were abolished when either VIPA^mut^ or the catalytically inactive 3C mutant (C172A) was expressed (*23*) (Fig. 1, C and D). Western blot analysis further confirmed that 3C protease efficiently cleaved full-length (FL) VIPA to release its N-terminal fragment (VIPA^NT^) (Fig. 1E).

We next assessed the impact of VIPA on HAV infection in Huh7.5.1 cells (*24*). Immunofluorescence staining showed that heterologous expression of VIPA markedly inhibited the accumulation of viral VP1 protein (Fig. 1F) and significantly reduced HAV genomic RNA levels at 4-6 days post-infection (dpi) (Fig. 1G) in HAV-infected cells. Consistently, VIPA expression in HAV-infected cells triggered pyroptosis, characterized by progressive membrane swelling and rupture starting (Fig. 1H), coinciding with a sharp increase in LDH release (Fig. 1I). Together, these observations demonstrate that VIPA selectively induces pyroptosis in HAV-infected cells, leading to efficient elimination of the infected cells *in vitro*.

### Therapeutic efficacy of LNP-delivered VIPA mRNA in mice

To evaluate the *in vivo* therapeutic potential of VIPA, we first determined the expression and safety profile of VIPA-encoding mRNAs in mice. Following intravenous (i.v.) injection of VIPA mRNA-LNP, robust hepatic expression of VIPA protein was observed in mouse liver at 24 h post-injection (fig. S1A). Meanwhile, no obvious changes were detected in liver function tests or histopathological examinations (fig. S1, B and C), supporting a favorable safety profile. We next evaluated the antiviral efficacy of VIPA in an established *Ifnar^-/-^* mouse model of HAV infection (Fig. 2A) (*25*). All placebo-treated mice exhibited sustained viral shedding in feces up to 18 dpi, whereas VIPA-treated mice showed a marked reduction in fecal HAV RNA levels, approximately 100-fold lower than those in placebo controls (Fig. 2B). Consistently, hepatic HAV RNA loads were significantly reduced (Fig. 2C), and *in situ* RNA scope analysis confirmed the near-complete absence of HAV RNA-positive hepatocytes (Fig. 2D). More importantly, serum biochemical assays showed that VIPA-treated mice predominantly maintained ALT and AST levels within the physiological range throughout the observation period (Fig. 2E), whereas severe liver dysfunction was observed in all placebo-treated mice, as previously described (*25*). Histopathological examination revealed multifocal inflammatory infiltrates surrounding necrotic or apoptotic hepatocytes in the placebo-treated animals, while no overt pathological abnormalities were observed in the livers of VIPA-treated mice (Fig. 2F).

**Fig. 2.**
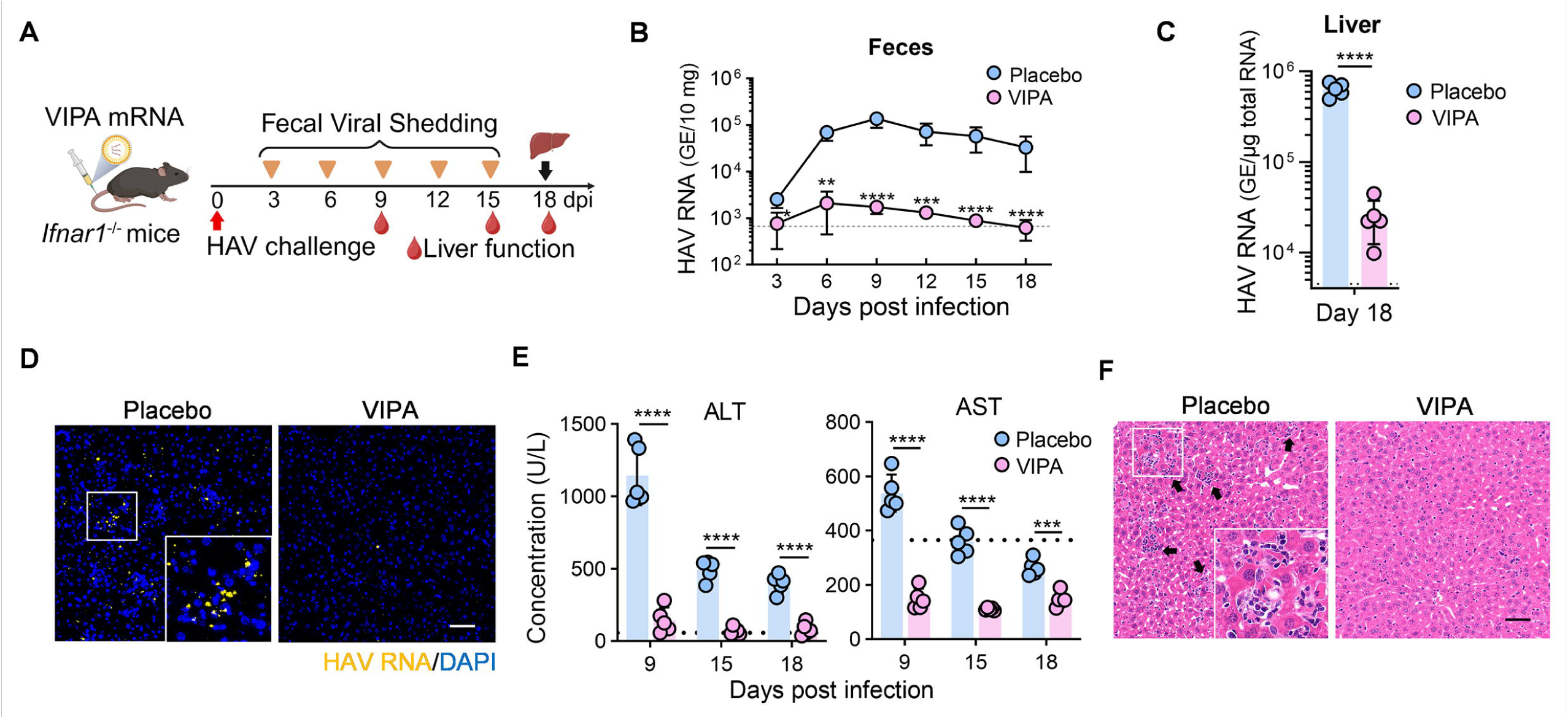
Therapeutic efficacy of VIPA mRNA in mice. **(A)** Schematic of the *in vivo* assessment of VIPA antiviral efficacy (created with BioRender.com). *Ifnar*^−/−^ mice (n = 5) were infected with HAV vRNA formulated in lipid nanoparticles (LNP) and treated with VIPA mRNA-LNP *via* tail vein injection. The red arrow indicates the time point of HAV challenge, yellow triangles indicate fecal sample collection, red droplets indicate blood sampling for liver function analysis, and the black arrow marks the endpoint when mice were euthanized for tissue analyses. **(B)** Time course of HAV shedding in feces from mice treated with VIPA mRNA-LNP or placebo at 3, 6, 9, 12, 15, and 18 dpi. **(C)** Quantification of HAV RNA copies in liver tissues at 18 dpi by RT-qPCR. **(D)** RNAscope *in situ* hybridization detecting HAV vRNA in liver sections. Yellow puncta indicate HAV RNA signals. Scale bar, 25 μm. **(E)** Serum alanine aminotransferase (ALT) and aspartate aminotransferase (AST) levels measured at 9, 15, and 18 dpi. Dashed lines indicate the safety threshold. **(F)** Representative hematoxylin-eosin (H&E)-stained liver sections at 18 dpi. Typical HAV-associated inflammatory foci observed in placebo-treated mice are highlighted in enlarged views. Scale bar, 25 μm. Data are presented as mean ± SD.

Additionally, comparable antiviral and hepatoprotective effects were observed in another established mouse model (*26*) infected with the HAV strain HM175/18f (fig. S2, A to C). Collectively, these findings demonstrate that LNP-delivered VIPA mRNA therapy efficiently suppresses HAV replication and fecal viral shedding across multiple viral strains, and alleviates HAV-associated long-term liver injury *in vivo*.

### VIPA mRNA promotes coordinated antiviral responses in liver immune cells, enhancing T cell function

To delineate the mechanism of VIPA-mediated protection, we first examined hepatic cytokine profiles at 9 dpi. VIPA-treated mice showed no elevation of cytokine storm-associated cytokines (IL-6, IL-1β, TNF-α) but displayed selective upregulation of specific chemokines (fig. S3A), indicative of localized and regulated inflammation rather than systemic immune activation. Bulk transcriptomic analysis of liver tissue revealed significant enrichment in pathways related to pathogen recognition, antigen presentation, interferon (IFN) production and response, and T cell activation pathways (Fig. 3A). These findings collectively suggest that VIPA-induced pyroptosis in hepatocytes initiates a coordinated antiviral immune program.

**Fig. 3.**
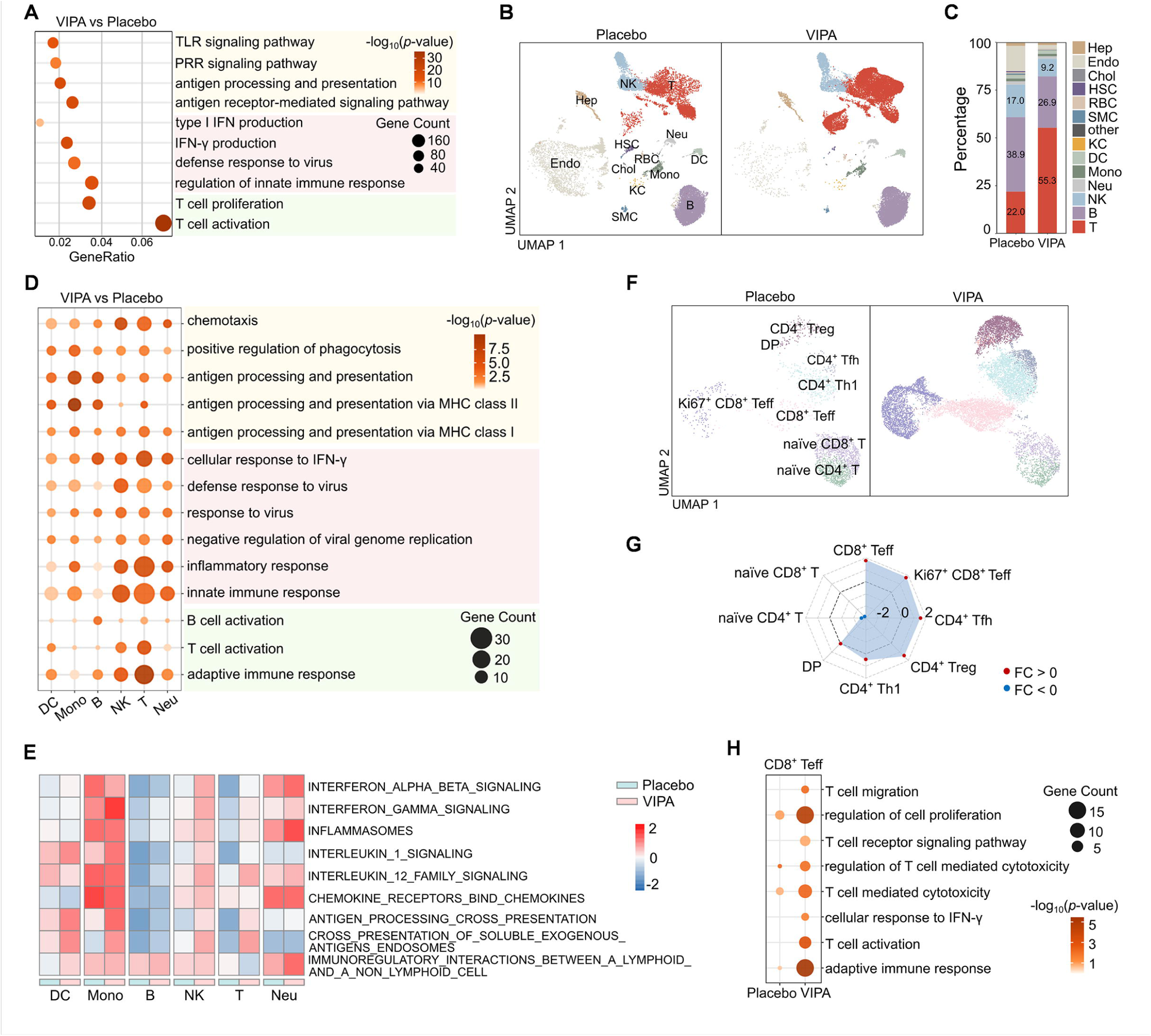
VIPA mRNA therapy triggers antiviral immune response in uninfected liver immune cells. **(A)** Bubble plot of antiviral signaling pathways upregulated in liver tissues of HAV-infected mice treated with VIPA vs placebo at 9 dpi (bulk RNA-seq, n = 5). **(B)** UMAP of major cell populations identified by scRNA-seq in mouse liver, comparing VIPA and Placebo groups (9 dpi, n = 3). **(C)** Proportions of major intrahepatic cell types in Placebo and VIPA groups (bar plots). **(D)** GO enrichment of upregulated genes in intrahepatic immune cell populations from mouse liver (VIPA vs Placebo). **(E)** Heatmap of GSVA scores for antiviral REACTOME gene sets across cell types and treatment groups (VIPA vs Placebo). **(F)** UMAP visualization of T cell subclusters comparing VIPA and Placebo groups from scRNA-seq data. **(G)** Radar plot of fold changes in major T cell subtypes between VIPA and Placebo groups. **(H)** Bubble plot of enriched T cell activation signaling pathways in CD8^+^ T cells from VIPA and Placebo group versus mock-treated mice (scRNA-seq, upregulated genes).

Single-cell RNA-seq of mouse liver revealed 13 distinct immune and parenchymal cell populations (Fig. 3B and fig. S3, B and C). VIPA mRNA therapy increased the proportion of T cells with higher gene-signature scores (*27*) compared to placebo, while concomitantly reducing the proportions of B cells, natural killer (NK) cells, monocytes (Mono), dendritic cells (DCs), endothelial cells (Endo), and hepatocytes (Hep) (Fig. 3C and fig. S3E). Gene Ontology (GO) enrichment analysis demonstrated that major immune cell populations, including T cells, NK cells, and neutrophils (Neu), exhibited robust antiviral activation signatures following VIPA mRNA treatment (Fig. 3D). These signatures encompassed innate antiviral responses, responsiveness to IFN-γ, suppression of viral replication, and enhanced chemotaxis and adaptive immune response. A similar antiviral response profile was also observed in other immune cell types, including B cells, Mono, and DCs. Gene Set Variation Analysis (GSVA) (*28*) further indicated that VIPA activated key signaling pathways, including IFN-α/β/γ, IL-1/12, and antigen processing pathways in most immune cells (Fig. 3E).

Sub-clustering analysis further revealed an expansion of CD8^+^ effector and CD4^+^ helper T cell subsets, accompanied by a reduction in naïve populations (Fig. 3, F and G). Differential expression analyses showed upregulation of genes associated with cytokine secretion in CD4^+^ T cells and migration and cytotoxicity in CD8^+^ T cells (fig. S3, I and J). GO enrichment analysis highlighted enhanced effector functions, cytokine production, and antiviral responses in T cells following VIPA treatment (Fig. 3H and fig. S3K).

Together, these data demonstrate that VIPA mRNA therapy not only kill infected hepatocytes but also reprograms uninfected hepatic immune cells into an antiviral alert state, enhancing CD8^+^ effector T cell function to promote HAV clearance (*29*).

### Design and characterization of VIPA against ZIKV and SARS-CoV-2

To further validate the concept of VIP strategy with other viruses with global impact, ZIKV-responsive VIPA was designed and constructed accordingly (fig. S4A). *In vitro* assays confirmed that ZIKV VIPA was specifically cleaved by viral NS2B3 protease (*30*), leading to robust pyroptosis and LDH release in HEK293T cells (fig. S4, B to E).

As expected, expression of ZIKV VIPA triggered robust pyroptotic cell death, and markedly reduced viral RNA replication in ZIKV-infected HeLa cells (Fig. S4, F to I). *In vivo* delivery of ZIKV VIPA mRNA-LNP substantially reduced viral RNA loads in liver (fig. S4, J to K); Consistently, liver sections displayed characteristic pyroptotic morphology in VIPA-treated mice (fig. S4L) (*31*). VIPA treatment also enhanced antiviral immunity, evidenced by early activation of type I interferon signaling and elevated serum IFN-β levels. (fig. S4M). Similarly, a SARS-CoV-2-responsive VIPA, incorporating the viral main protease NSP5 cleavage motif (*32*), exhibited potent antiviral activity by triggering pyroptotic cell death (Fig. 4, A and B and fig. S5, A to F). Collectively, these results demonstrate that the VIP approach can be adapted to distinct viral species, enabling selective elimination of infected cells and amplification of host antiviral responses.

**Fig. 4.**
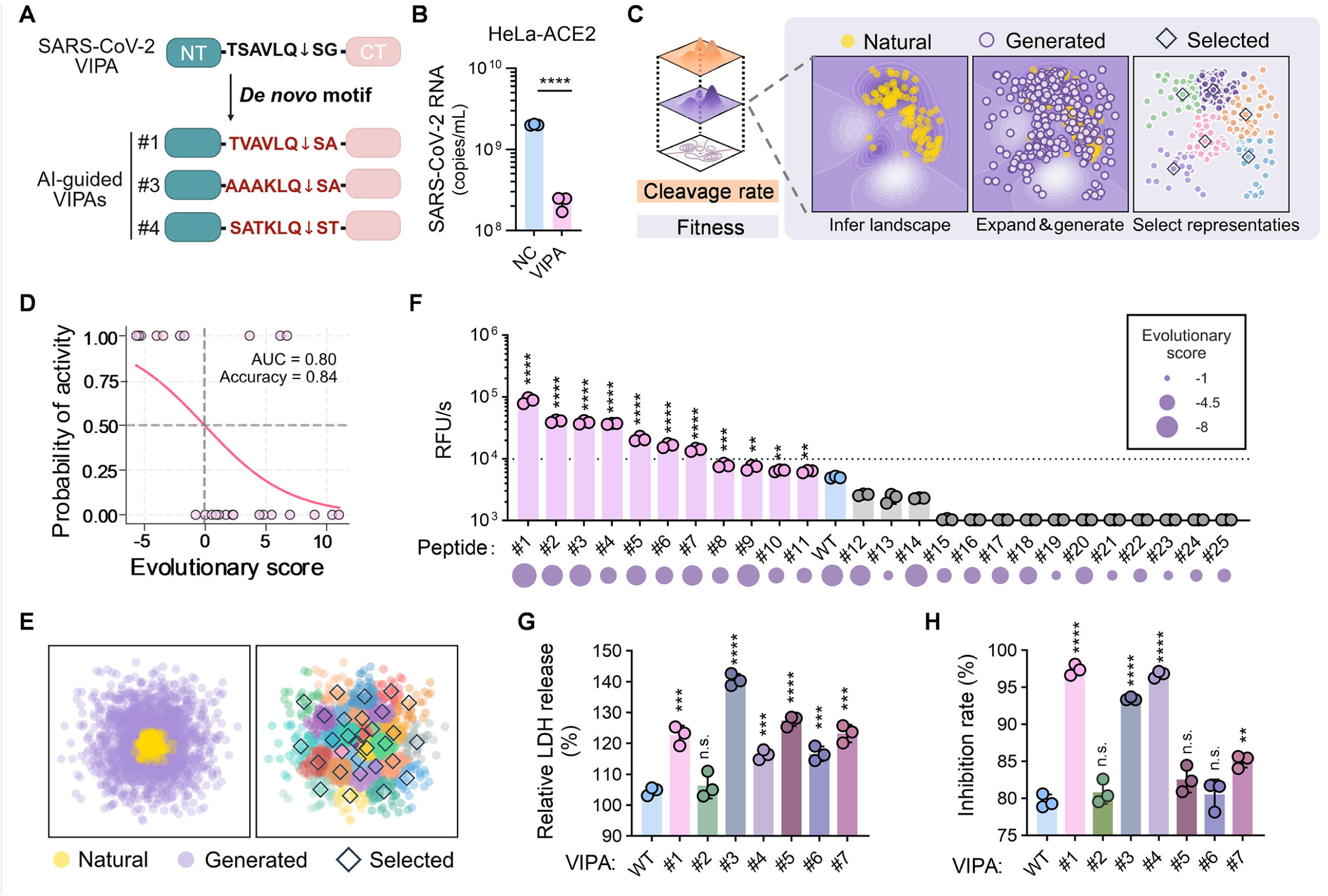
AI-guided VIPA design against SARS-CoV-2. **(A)** Schematic of original and AI-guided VIPA constructs targeting SARS-CoV-2, created with BioRender.com. **(B)** HeLa-ACE2 cells (n = 3) were transfected with VIPA and infected with SARS-CoV-2. Viral RNA replication was quantified by RT-qPCR. **(C)** Framework of the AI-guided VIPA exploration workflow. An unsupervised VAE model learns the evolutionary constraints from natural sequences (yellow); generates candidates (purple), which are clustered; centroid-proximal representatives (diamonds) are selected for experimental assays. Fitness (purple) and cleavage rate (orange) landscapes are related but distinct; The 2D plane depicts the sequence space, surfaces depict each landscape. **(D)** The performance of the evolutionary score in predicting cleavage outcomes on an independent test set (n = 25) was evaluated, yielding an F1 score of 0.78. **(E)** Left: Qualitative visualization of the distribution of natural (yellow; n = 261) and AI-generated sequences (purple; CD-HIT-deduplicated and score-filtered, n = 3,305). Right: K-Means clusters (colors) and selected candidates (diamonds). **(F)** Cleavage kinetics (ΔRFU/s) of representative AI-generated peptide candidates measured in the FRET assay, showing variable susceptibility to NSP5-mediated cleavage. **(G)** LDH release assay in HEK293T cells (n = 3) co-expressing NSP5 and each AI-guided VIPA candidate, identifying variants with enhanced pyroptotic activity. **(H)** Antiviral activity of each AI-guided VIPA candidate in HeLa-ACE2 cells (n = 3), quantified as inhibition of SARS-CoV-2 replication using viral RNA levels. Data are presented as mean ± SD.

### AI-guided VIPA generation and validation

Given that viral genomes including the protease cleavage motifs are continuously evolving (*33*), we sought to harness AI-based computational pipeline to enhance the antiviral activity and mutation tolerance of VIPAs. Herein, we proposed a generation-sampling co-design framework -- "learn the known space, explore unobserved regions, representative sampling, and experimental validation" -- to guide data-driven exploration of viral protease substrate landscape (Fig. 4C and fig. S6A).

Specifically, we trained an unsupervised evolutionary model of variant effect (EVE), a variational autoencoder (VAE)-based model (*34*), using 261 known natural SARS-CoV-2 NSP5 cleavage motifs spanning positions P6-P2′ (*35*) curated from the NCBI Virus database. Model performance was validated on an independent test set of 25 sequences with experimentally measured cleavage outcomes, yielding the model’s evolutionary score that effectively discriminated cleavable from non-cleavable motifs (ROC-AUC = 0.80; accuracy = 0.84) (Fig. 4D and fig. S6B).

Then, 100,000 candidate sequences were generated and embedded based on pairwise sequence similarities using principal coordinates analysis (PCoA): the generated set covered the region occupied by natural sequences and extended into previously unobserved peripheral regions (Fig. 4E and fig. S6E). Using a **cluster-based representative sampling**, we selected 30 diverse motifs for experimental validation. (Fig. 4E).

To experimentally evaluate protease responsiveness, we established an *in vitro* FRET-based assay to quantify NSP5 cleavage kinetics for each candidate peptide (fig. S6G). Among selected octapeptides, seven of the 25 successfully synthesized candidates exhibited more than a twofold increase in cleavage rate compared with the wild-type (WT) substrate (Fig. 4F). Consistent with this, the evolutionary score correlated with experimentally measured cleavage efficiency across 25 candidates (Spearman’s rank, p < 0.001).

These top candidates were individually incorporated into the VIPA scaffold to generate corresponding variants (VIPA#1 to 7). Functional assays in NSP5-expressing HEK293T cells showed that six of the seven AI-guided VIPAs triggered stronger LDH release than the WT VIPA (Fig. 4G). Moreover, in SARS-CoV-2-infected HeLa cells, three variants (VIPA#1, #3, and #4) exhibited significantly enhanced antiviral potency (inhibition rate > 90%) relative to the WT construct (Fig. 4H). Collectively, these AI-guided, evolution-informed VIPAs (Fig. 4A) represents potent candidates for further development.

## Discussion

VIPA represents a conceptual advance in the design of precision antiviral therapeutics, by rewiring non-specific pyroptosis host defense into a viral protease-responsive antiviral response, thereby selectively eliminating infected cells while sparing uninfected ones. A key feature of VIPA lines in its exceptional specificity and favorable safety profile. Previous approaches to modulate GSDM activity, such as small molecule inhibitors, antibodies, or peptide blockers (*11*, *12*, *36–38*), lack cell-type or tissue specificity and pose a risk of unintended pyroptosis in bystander cells. In contrast, VIPA restricts GSDM activation to virally infected cells *via* recognition and cleavage of viral proteases, ensuring that pyroptosis is strictly confined within these cells. More importantly, delivery of VIPA mRNA conferred robust antiviral protection without detectable hepatotoxicity or systemic cytokine storm across multiple animal models, further highlighting its favorable safety profile (fig. S1, B and C and fig. S3A).

Theoretically, VIPA can be flexibly adapted to target a broad range of viruses that encode proteases. In our study, VIPAs directed against HAV, ZIKV and SARS-CoV-2 exhibited promising antiviral effects in multiple *in vitro* and *in vivo* models. Furthermore, the modular design of VIPA also allows for the tandem incorporation of multiple viral protease motifs. Such "poly-specific" VIPA constructs can respond to distinct protease activities from multiple or co-infecting viruses, thereby broadening antiviral coverage and reducing the likelihood of viral escape through protease mutations. This inherent flexibility positions VIPA as a modular antiviral platform that can be rapidly customized to counter emerging "X" viral threats. Using an unsupervised generative evolutionary model (*39–41*), we explored and expanded the natural protease recognition space, identifying optimized VIPAs with significantly enhanced cleavage kinetics and antiviral efficacy. Theoretically, these AI-guided VIPAs would maintain antiviral efficacy even when viral variants acquire mutation at their protease cleavage sites. The same computational framework can be applied to optimize VIPAs targeting other viruses.

Compared with other mRNA-based antiviral therapies under development (*42–45*), VIPA not only induces a controlled, cell-intrinsic suicide response specifically in infected cells, but also alerts neighboring uninfected cells. RNA-seq data from VIPA-treated mice demonstrated that most immune cells, including DCs, Mono, neutrophils, NK cells, and T cells, were induced into an antiviral state, with concurrent activation of both innate and adaptive immune responses (Fig. 3 and fig. S3). Notably, VIPA-mediated pyroptosis reshapes the immune microenvironment, driving the activation of IFN-related antiviral pathways and facilitating the expansion and functional activation of effector CD8^+^ and CD4^+^ T cells (Fig. 3 and fig. S3), which are essential for effective virus clearance (*29*, *46*). Thus, VIPA-induced release of DAMPs and PAMPs may act as a localized adjuvant, bridging innate and adaptive immunity to facilitate viral clearance without causing systemic inflammation. This mechanism fundamentally differs from classical antiviral strategies, thereby establishing VIPA as a "kill-and-alert" paradigm.

Overall, our study establishes a framework for designing protease-activated pyroptotic switches. Using GSMD as a scaffold, other therapeutic mRNAs targeting different viruses as well as pyroptosis -related diseases are currently under development for further clinical translation.

## Data Availability

All data supporting the findings of this study are available from the corresponding author upon reasonable request. Upon acceptance, all relevant data will be deposited in a publicly accessible repository, and any persistent URLs or accession numbers will be provided at that time. Additional raw images and analysis scripts have been included as Supplementary Material.

## Code Availability

All custom code implementing the learn-expand-sample workflow is available at https://github.com/wuaipinglab/eve-lexs-nsp5 under the Apache License 2.0.

## Acknowledgements

This work was supported by grants from the National Key Research and Development Program of China (2021YFC2302400, 2022YFC2303700), National Natural Science Foundation of China (32500551, 32130005), Major Project of Guangzhou National Laboratory (GZNL2024A01015), and State Key Laboratory of Pathogen and Biosecurity (SKLPBS2405). The data analysis process has been supported by the High-throughput Sequencing and High-performance Computing Platform of the Suzhou Institute of Systems Medicine, Chinese Academy of Medical Sciences & Peking Union Medical College.

## Author Contribution Statement

C.-F.Q., L.L., and Y.-J.H. conceived the project. C.-F.Q., Y.-J.H., and A-P.W. supervised the study. C.-F.Q., L.L., and Y.-J.H. wrote the manuscript with input from all authors. L.L., Y.-J.H., and X.-L.Y. designed the experiments. L.L. performed the mRNA-LNP preparation, biochemical assays, cell-based experiments and animal studies, assisted by X.-L.Y., Y.-J.H., J.L., R.-R.Z., L.Lv., X.-Y. H., T.-S.C., Q.Y. and H.Z.. H.-Y.W., H.-Y.Z. and H.D. conducted the AI-related and statistical analyses with support from L.L.. C.-F.Q., L.L., Y.-J.H., X.-L.Y., Y.F. and A.W. analyzed the data. L.L., X.-L.Y., and Y.-Y.L. performed RNA-seq analyses.

## Conflicts of Interest Statement

The authors have filed patents related to this manuscript.

