## Supplementary materials for "Viral protease-Initiated Pyroptosis Activator mRNA therapy as a Universal Antiviral Strategy"

### Methods

### Cell culture

Human cell lines including HeLa (cervical carcinoma, ATCC CCL-2), HeLa-ACE2, HEK293T (embryonic kidney, ATCC CRL-11268), and Huh7.5.1 (hepatoma, Procell CL-1096) were cultured in Dulbecco’s Modified Eagle’s Medium (DMEM, Gibco 11995065) supplemented with 10% fetal bovine serum (FBS, Gibco A5670701) and 1% penicillin-streptomycin (P/S, Gibco 15140122) at 37 °C in a humidified incubator with 5% CO₂. Animal cell lines including BHK-21 (baby hamster kidney, ATCC CCL-10) and Vero (African green monkey kidney, ATCC CCL-81) were maintained under the same conditions. Mosquito cell line C6/36 (Aedes albopictus, ATCC CRL-1660) was cultured in RPMI-1640 medium (Gibco 11875093) supplemented with 10% FBS and 1% P/S at 28 °C without CO₂. All cell lines were routinely tested and confirmed negative for mycoplasma contamination before use.

### Virus propagation and quantification

HAV strain HM175/18f was propagated in Huh7.5.1 cells. The viral RNA (vRNA) of HAV strain HM175-mp4 was synthesized according to the procedure described in the mRNA synthesis section. Both viral preparations were quantified by RT-qPCR using specific primers (Forward: 5'-GGTAGGCTACGGGTGAAAC-3'; Reverse: 5'-AACAACTCACCAATATCCGC-3'). Zika strain MR766 was amplified in C6/36 cells, and viral titers were determined by RT-qPCR and plaque assay on BHK cells. (Primers F: 5'-CCgCTgCCCAACACAAg-3', R: 5'-CCACTAACgTTCTTTTgCAgACAT-3', Probe: 5'-FAM-gCCTACCTTgACAAgCAATCAgACACTCAA-BHQ1-3'). SARS-CoV-2 strain BetaCoV/Beijing/IME-BJ01/2020 (131) (*1*) was propagated in Vero cells and quantified by RT-qPCR (Primers F: 5’-TCCTGGTGATTCTTCTTCAGGT-3’, R: 5’-TCTGAGAGAGGGTCAAGTGC-3’, Probe: 5’-FAM-AGCTGCAGCAC CAGCTGTCCA-BHQ1-3’).

All experiments involving infectious HAV, ZIKV, and SARS-CoV-2 were performed in the appropriate biosafety containment facilities at Academy of Military Medical Sciences (AMMS).

### Animals

All animal experiments were conducted in full compliance with the Regulations for the Administration of Laboratory Animals and the Laboratory Animal-Requirements of Environment and Housing Facilities of China. Experimental procedures were reviewed and approved by the Institutional Animal Care and Use Committee of the AMMS (protocol numbers: IACUC-DWZX-2025-A010 and IACUC-DWZX-2025-A002). Every effort was made to ensure the welfare of the animals, including careful attention to housing environment, handling practices, and the reduction of pain or distress throughout the study.

C57BL/6J mice were purchased from Beijing Vital River Laboratory Animal Technology Co., Ltd. *Ifnar1*^-/-^ mice were bred by Suzhou Cyagen Biosciences Inc. Animals were randomly assigned to experimental and control groups.

### VIPA construction

VIPA was constructed by fusing the N-terminal pore-forming domain of GSDMD (residues 1-258), a viral protease recognition motif, and the C-terminal inhibitory domain (residues 276-484). To prevent unintended activation of endogenous GSDMD-associated pathways and to avoid potential loss of function caused by off-target viral protease cleavage, six residues (Q29A, D87A, Q193A, R249A, L290A, and R291A) were mutated (*2*–*6*). For virus-specific targeting, the cleavage motifs were derived from the cognate proteases of HAV (LRTQ↓SF), ZIKV (KTGKR↓SG), and SARS-CoV-2 (TSAVLQ↓SG), enabling selective activation of VIPA by the corresponding viral enzymes. In the control construct (VIPA^mut^), the protease-recognition linker was replaced with a scrambled sequence to abolish protease responsiveness.

### mRNA synthesis

mRNAs encoding VIPAs were synthesized in vitro using the T7 High Yield RNA Transcription Kit (Vazyme, TR101-01) from linearized plasmid templates (GenScript) containing the full coding sequences flanked by 5′ and 3′ untranslated regions. Each transcript included a polyA tail of more than 100 nucleotides.

The full-length cDNA of HAV HM175-mp4 (GenBank accession no. KX343018.1) was synthesized by Tsingke Biotech and cloned into the pGEM3 vector to generate the recombinant plasmid pGEM3-HM175-mp4 (*7*). Viral RNA (vRNA) was subsequently transcribed in vitro from this plasmid using the RiboMAX™ Large Scale RNA Production System-T7 (Promega) to produce capped HAV vRNA for downstream experiments. All mRNA preparations were analyzed using an Agilent 2100 Bioanalyzer to confirm integrity and purity, with all samples exhibiting >95% purity.

### Lipid-nanoparticle encapsulation of the mRNA

The formulation of lipid nanoparticles (LNP) was executed based on a well-defined protocol as previously described (*1*). The process commenced with the preparation of a lipid mixture, which included 1,2-distearoyl-sn-glycero-3-phosphocholine (DSPC), cholesterol, and a PEGylated lipid species, dissolved in ethanol at molar ratios of 50:10:38.5:1.5. This lipid solution was then combined with a 20 mM citrate buffer (pH 4.0, Teknova, Q2444) containing mRNA at a 1:2 volume ratio, using the NanoAssemblr Ignite™ platform (Cytiva). The resulting suspension was subsequently dialyzed against PBS (pH 7.4, Gibco, 70011044) using a 20 kDa molecular weight cutoff membrane, followed by ultrafiltration to concentrate the LNPs. The final product was filtered through a 0.22 µm membrane and stored at 4°C. The final preparation was passed through a 0.22 µm filter and stored at 4 °C. Each formulation underwent comprehensive characterization, including measurements of particle size, size distribution, RNA content, and encapsulation efficiency.

### Plasmid transfection and fluorescence imaging

HEK293T cells were seeded in confocal dishes at a density of 2×10⁵ cells/well. After 12 h, cells were transfected with pcDNA3.1-VIPA, with or without the corresponding viral protease, using Lipofectamine 3000 (Thermo Fisher Scientific, L3000015). Six hours later, Annexin A5-FITC (Vazyme, A211-01) was added to the culture medium and incubated for 5 min. Fluorescence images were acquired using a Zeiss LSM980 confocal laser scanning microscope equipped with an AiryScan super-resolution module.

### Lactate dehydrogenase release and ATP measurement

HEK293T cells were seeded in 12-well plates at a density of 2×10⁵ cells/well and transfected with pcDNA3.1-VIPA, with or without the corresponding viral protease, using Lipofectamine 3000 (Thermo Fisher Scientific, L3000015). After 12 h, cytotoxicity was assessed by measuring lactate dehydrogenase (LDH) release using the CytoTox 96 Non-Radioactive Cytotoxicity Assay (Promega), and cell viability was evaluated using the CellTiter-Glo Luminescent Cell Viability Assay (Promega). For viral infection experiments, LDH release was measured at the indicated time points using the same assay protocol.

### HAV infection and animal experiments

*Ifnar*^-/-^ C57BL/6 female mice (7 weeks old) were intravenously inoculated with either HAV HM175-mp4 vRNA encapsulated in LNPs or infectious HAV HM175/18f strains. VIPA mRNA-LNPs were administered intravenously at a dose of 5 μg per mouse per day. Fecal and serum samples were collected at predetermined time points to quantify viral RNA loads and measure serum ALT levels. Liver tissues were harvested for RNA extraction, H&E staining, immunofluorescence, RNAscope, bulk RNA-seq, and single-cell RNA-seq analyses. Samples designated for RNA extraction were preserved in TRIzol reagent, while those for histological analysis were fixed in 4% paraformaldehyde.

### ZIKV infection and animal experiments

Four-week-old female C57BL/6 mice were pretreated with anti-IFNAR1 monoclonal antibody (0.5 mg per mouse, intravenously) one day before ZIKV infection to transiently block type I interferon signaling. Mice were then **injected intraperitoneally** with ZIKV (1 × 10^7^ PFU per mouse) on the following day.

ZIKV VIPA mRNA-LNPs were administered intravenously at 5 μg/mouse/day, beginning on the day of infection. Serum samples were collected at 2 and 5 dpi for quantification of IFN-α, IFN-β, and IFN-γ.

At designated time points, liver tissues were harvested for RNA extraction and histopathological examination. Samples for RNA analysis were preserved in TRIzol reagent (Invitrogen), whereas those for histology were fixed in 4% paraformaldehyde and processed for H&E staining.

### **Blood biochemistry and cytokine analysis**

Blood samples were collected from HAV-infected mice at 9, 15, and 18 dpi. Samples were allowed to clot at room temperature for 1 h, and serum was separated by centrifugation at 6,000 × g for 10 min.

Serum ALT and AST levels were measured using an automatic biochemistry apparatus (Hitachi3100, HITACHI) corresponding commercial assay kits conducted by Beijing Biocytogen Co., Ltd. For cytokine profiling, concentrations of 36 cytokines were quantified using the Mouse ProcartaPlex 36-Plex Panel (Invitrogen, EPX360-26092-901) on a Luminex 200 platform, following the manufacturer’s instructions, with measurements performed by Shanghai Laizee Biotech Co., Ltd.

### Real-time quantitative PCR

Total RNA was extracted from cells and tissues using TRIzol reagent (Invitrogen, USA, 15596026) following the manufacturer’s protocol. For fecal samples, RNA was isolated after homogenization in PBS (pH 7.4) using the qEx-DNA/RNA Kit (TIANLONG, China, T335). HAV genomic RNA was quantified with the One Step TB Green PrimeScript™ PLUS RT-PCR Kit (Takara, Japan, RR096A), using primers described previously. ZIKV and SARS-CoV-2 genomic RNAs were quantified using the One Step PrimeScript RT-PCR Kit (Takara, Japan, RR064A) with virus-specific primer pairs. All reactions were performed on a Roche LightCycler^®^ 480 Real-Time PCR System. Absolute RNA copy numbers were determined using a standard curve generated from synthetic HAV RNA transcripts.

### **Bulk RNA-seq library preparation and sequencing**

Total RNA was extracted from mouse liver tissues collected at 9 days post-infection with HAV using TRIzol reagent (Invitrogen) according to the manufacturer’s instructions. Poly(A)^+^ mRNA was purified from total RNA using Oligo(dT)-attached magnetic beads and subsequently fragmented into short fragments. First-strand cDNA was synthesized using random hexamer primers, followed by second-strand synthesis. For strand-specific libraries, dUTP was incorporated in place of dTTP during second-strand synthesis. The resulting double-stranded cDNA was subjected to end repair, A-tailing, adaptor ligation, size selection, PCR amplification, and purification.

Library quality was assessed using a Qubit fluorometer (Thermo Fisher Scientific) for concentration measurement and an Agilent 2100 Bioanalyzer for fragment size distribution. Quantified libraries were pooled according to the desired sequencing depth and sequenced on an Illumina NovaSeq 6000 platform using paired-end (2 × 150 bp) sequencing-by-synthesis chemistry.

### RNA-seq data processing and analysis

Raw sequencing reads were processed using fastp (*8*) (v0.23.4) to trim adapter sequences, remove low-quality reads, and discard reads containing more than 10% ambiguous bases (Ns). Clean reads were aligned to the Mus musculus reference genome (GRCm38) using HISAT2 (*9*) (v2.2.1) with corresponding gene model annotations to improve splice junction mapping. Gene-level read counts were quantified using featureCounts (*10*) (v2.0.6) and normalized to fragments per kilobase of exon per million mapped reads (FPKM). Differential expression analysis was performed using DESeq2 (*11*) (v1.42.0) for datasets with biological replicates, or edgeR (*12*) (v4.0.16) when replicates were unavailable. The Benjamini–Hochberg method was applied to control the false discovery rate (FDR), and genes with adjusted P ≤ 0.05 and |log₂(fold change)| ≥ 1 were considered significantly differentially expressed. Functional enrichment analyses, including Gene Ontology (GO) and KEGG pathway analyses, were conducted using clusterProfiler (*13*) (v4.8.1).

### Single-cell RNA-seq library preparation and sequencing

Liver tissues from mice at 9 days post HAV infection were immediately transferred to ice-cold RNase-free PBS (without Ca²⁺ and Mg²⁺) and minced into approximately 0.5 mm^3^ pieces. Tissues were washed thoroughly with PBS to remove residual blood and lipids, then enzymatically digested in 0.35% collagenase IV, 2 mg/mL papain, and 120 U/ml DNase I at 37 °C for 20 min with gentle agitation (100 rpm).

Digestion was quenched with PBS containing 10% FBS, and the suspension was gently dissociated by pipetting and filtered sequentially through 70 µm and 30 µm strainers. Cells were pelleted by centrifugation at 300 × g for 5 min at 4 °C and resuspended in PBS with 0.04% BSA. Red blood cells were lysed using 1× RBC Lysis Buffer (MACS 130-094-183) for 5 min on ice. Dead cells were removed using Dead Cell Removal MicroBeads (MACS 130-090-101) according to the manufacturer’s instructions. The final cell pellet was washed twice with PBS + 0.04% BSA and resuspended in 50 µl of the same buffer. Cell viability (>85%) was assessed by trypan blue exclusion, and cell concentration was adjusted to 700-1200 cells/µl for downstream library preparation.

Single-cell suspensions were loaded onto the 10x Chromium instrument following the manufacturer’s instructions (Chromium Single-Cell 3’ kit, V3), with a target capture of 10,000 cells per sample. Reverse transcription, cDNA amplification, and library construction were performed according to the kit protocol. Libraries were sequenced on a NovaSeq 6000 platform (Illumina) using paired-end 150 bp reads, with a sequencing depth of ~20,000 reads per cell. All library preparation and sequencing services were provided by LC-Bio Technology Co., Ltd. (Hangzhou, China).

### Single-cell RNA-seq data processing and analysis

Raw sequencing data were demultiplexed and converted into FASTQ format using Illumina bcl2fastq (version 2.20). The raw reads were processed with Cell Ranger (*14*) (version 8.0.1; 10x Genomics) for quality control, read alignment, and gene quantification against the Mus musculus reference genome (GRCm39, Ensembl release 105). In total, 89,382 cells were captured from nine mouse liver samples. The output matrices were imported into the Seurat R package (*15*) (version 5.2.1) for downstream analyses, including normalization, dimensionality reduction, clustering, and gene expression profiling. Potential doublets or empty droplets were removed using DoubletFinder (*16*) (version 2.0).

After quality control filtering, 74,225 high-quality cells were retained for further analysis. Filtering thresholds were set as follows: genes detected in at least three cells; each cell expressing ≥500 genes and ≥500 UMIs; and mitochondrial gene expression fraction <25%.

Filtered gene expression matrices were normalized using the LogNormalize method implemented in Seurat. Principal component analysis (PCA) was performed for dimensionality reduction, and the top 20 principal components were selected for subsequent clustering and visualization. Cell clusters were identified using the shared nearest neighbor (SNN) modularity optimization algorithm. Differentially expressed genes (DEGs) for each cluster were identified with the FindAllMarkers function in Seurat, using the bimod likelihood-ratio test as implemented in Seurat. Marker genes were defined as those expressed in ≥10% of cells within a cluster, with log fold change (logFC) ≥0.26 relative to all other cells.

### SARS-CoV-2 NSP5 cleavage motif dataset construction

SARS-CoV-2 ORF1ab amino acid sequences were retrieved from the NCBI Virus database (taxid: 2697049) using the following criteria: complete nucleotide sequences encoding the ORF1ab polyprotein, as of 2 May 2024, yielding 1,469,268 sequences. Multiple sequence alignment was performed using MAFFT (version 7) with the reference sequence YP_009724389.

Protease cleavage sites corresponding to NSP5 motifs (positions P6-P2′) were extracted from the aligned sequences. Sequences containing incomplete motifs, gaps, or ambiguous residues (N and X) were filtered out using custom Python scripts. Redundancy was removed by collapsing only exact‑identity P6-P2′ octapeptides; near‑identical motifs differing at any position were retained, yielding a final non-redundant set of 261 NSP5 cleavage motifs that served as the natural training set for downstream generative modeling.

### Learn-expand-sample computational workflow

We used EVE (evolutionary model of variant effect) (*17*) as the generative model with default hyperparameters (architecture, optimizer and learning rate, training epochs, etc.). The sequence re-weighting in MSA parameter θ was set to 0.01 (viral default). All training and inference were performed on a single NVIDIA V100 (16 GB).

To quantify evolutionary feasibility, we followed the EVEscape fitness component (*18*): for each position, the evolution index is the change in negative ELBO for the target sequence relative to the wild type; the sequence‑level evolutionary score (*E_s_*) is the sum over positions. Lower *E_s_* indicates closer proximity to the model‑learned evolutionarily feasible space. We used *E_s_* as an evolutionary prior and a quality‑control metric in downstream steps.

Because the posterior means and standard deviations of training samples in the EVE latent space are near 0 and 1, respectively (Extended Data Fig. 6c, d), we sampled latent variables from a standard normal prior and decoded with temperature 1 to promote diversity, generating 100,000 candidate sequences. After removing 105 sequences that duplicated the training or independent test sets, we computed *E_s_* for the remainder and obtained a pseudo‑probability of nsp5 cleavage using a logistic‑regression model. We then applied CD‑HIT (sequence identity threshold 0.65, all other parameters were left at their default values) for de‑redundancy clustering, combining the generated sequences with the training and independent test sets; after clustering, we retained only clusters composed solely of generated sequences.

Within each retained cluster, we selected the lowest‑*E_s_* sequence that was also predicted cleavable by logistic regression, yielding 3,305 candidates for further down‑sampling.

To obtain a diverse yet representative set under a limited sampling budget, we computed pairwise alignment similarities among candidates, defined distances as 1 − similarity, embedded the sequences to a 2D space via principal coordinates analysis (PCoA), and performed k‑means clustering in the embedded space (k = 30, matching the planned assay scale). We chose the sequence nearest each cluster centroid as the representative. Because initial representatives were enriched for the LQ motif at P2-P1, we applied a diversity‑promoting constraint: within each such cluster, we computed the 15th percentile (t) of centroid distances; if there existed a non‑LQ sequence with distance < t, we replaced the representative with the nearest non‑LQ sequence. After this constraint, the final 30 representatives included 18 non‑LQ sequences for wet‑lab evaluation.

### Kinetic screening of NSP5 substrate motifs

A high-throughput FRET-based assay was established to quantitatively assess the cleavage kinetics of 30 candidate octapeptides. Each peptide was synthesized with an N-terminal FAM fluorophore and a C-terminal Dabcyl quencher.

Recombinant NSP5 protease (Novoprotein, CR92) was diluted in assay buffer to a final working concentration of 5 ng/μL. Peptide substrates were freshly dissolved in DMSO (MCE, HY-Y0320C) and diluted in assay buffer to a final concentration of 30 μM immediately before use. Reactions were performed in black 96-well plates by mixing equal volumes of enzyme and substrate solutions, followed by incubation at 37 °C. Fluorescence was continuously monitored at 485/520 nm using a microplate reader (SpectraMax iD3, Molecular Devices). The initial reaction rates (ΔRFU/s) were determined from the linear phase of the kinetic curves after baseline correction (*19*). Substrates yielding reaction rates below 1 × 10³ RFU/s were considered poorly cleaved under these assay conditions.

### Statistics and reproducibility

Data were analyzed using GraphPad Prism (version 10) or custom Python scripts. EVE training and inference were implemented in PyTorch 2.4.0+cu124 using the reference EVE codebase pinned to GitHub commit 460d70efeeeded58bc69227a203540d68953ae88 (https://github.com/OATML-Markslab/EVE); logistic regression and k‑means were implemented using scikit‑learn 1.1.2; CD‑HIT version 4.8.1 was used for de‑redundancy. Unless otherwise indicated, values from experiments are presented as mean ± SD. Statistical significance between groups was determined using unpaired two-tailed t-tests. Significance levels are indicated as follows: n.s., not significant; **p* < 0.05; ***p* < 0.01; ****p* < 0.001; *****p* < 0.0001.

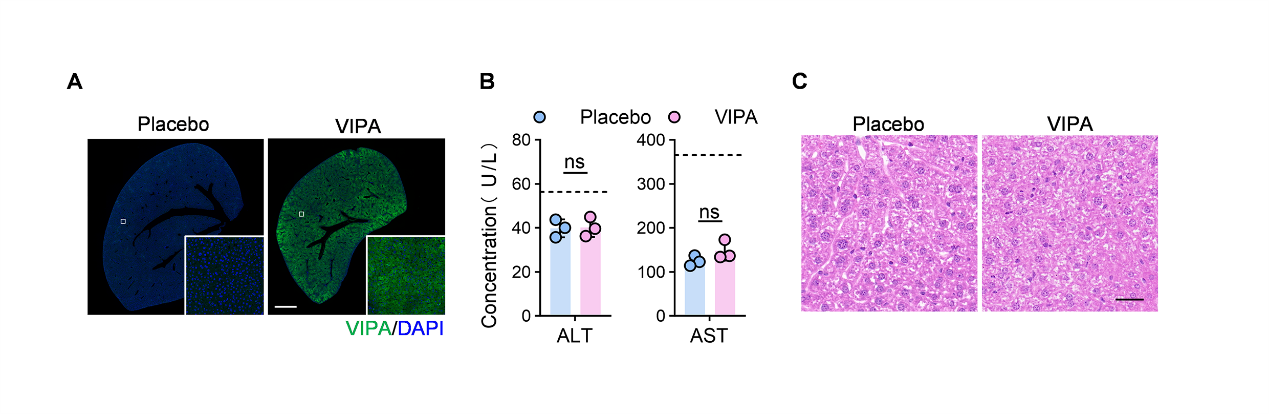

**fig. S1. Safety profile of VIPA-encoding mRNAs in mice.**

**(A)** Immunofluorescence showing VIPA protein expression (green) in mouse liver 24 h after injection of 5 μg VIPA mRNA-LNP. Scale bar, 1 cm.

**(B)** Serum ALT and AST levels were measured 24 h after injection of 5 μg VIPA mRNA-LNP. Dashed lines indicate the safety threshold.

**(C)** Representative H&E-stained liver sections 24 h after injection of 5 μg VIPA mRNA-LNP. Scale bar, 50 μm.

Data are presented as mean ± SD.

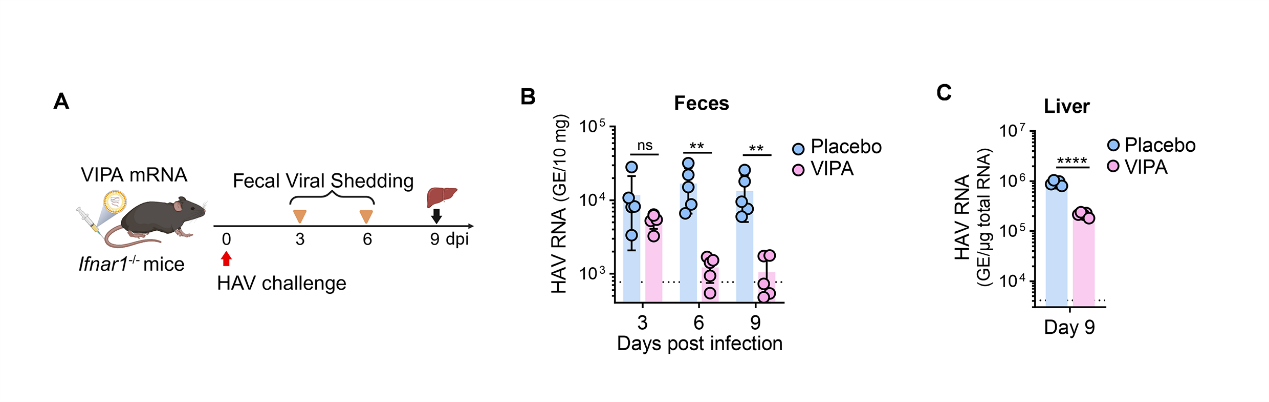

**fig. S2. Antiviral efficacy of VIPA mRNA in HAV-infected mice**

**(A)** Experimental design (created with BioRender.com). *Ifnar*^−/−^ mice (n = 5) were infected with HAV (HM175/18f) and subsequently treated with VIPA mRNA. Red arrow indicates time point of HAV challenge, yellow triangles indicate fecal sample collection, and black arrow denotes the experimental endpoint at which mice were euthanized for tissue analyses.

**(B)** Time course of HAV shedding in feces from mice in the VIPA or Placebo groups at 3, 6, and 9 dpi.

**(C)** Quantification of HAV RNA copies in liver tissues at 9 dpi by RT-qPCR.

Data are presented as mean ± SD.

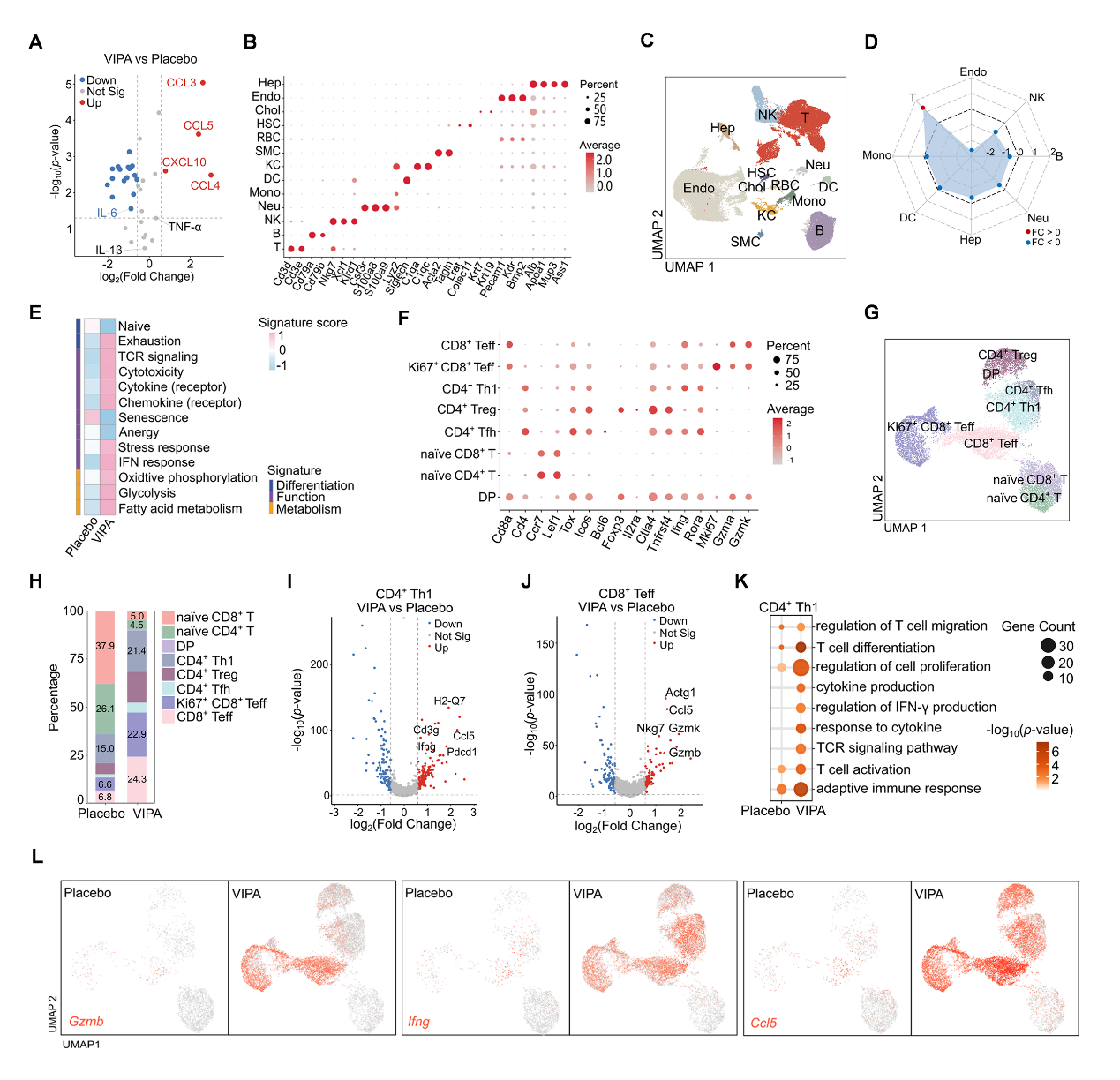
 **fig. S3.** Single-cell **RNA-seq (scRNA-seq) analysis of mouse liver following VIPA mRNA therapy.**

**(A)** Volcano plot showing differential cytokine levels in the livers of HAV-infected mice treated with VIPA mRNA compared with placebo controls at 9 dpi (n = 5). Upregulated cytokines are highlighted on the right, and downregulated cytokines on the left.

**(B)** Expression of representative marker genes defining each major cell type.

**(C)** UMAP visualization of scRNA-seq data depicting the compositional landscape of liver cell populations in HAV-infected mice at 9 dpi.

**(D)** Radar plot showing the relative proportions of major cell populations (>1% abundance) in VIPA versus Placebo-treated mice.

**(E)** Heat map illustrating expression of 13 curated gene signatures across all T cell clusters. Heat map was generated based on the scaled gene signature scores.

**(F)** Expression of canonical marker genes for distinct T-cell subsets.

**(G)** UMAP visualization of T-cell subclusters identified from scRNA-seq data.

**(H)** Bar plots showing the proportional changes in T-cell subsets between groups.

**(I and J)** Volcano plots of DEGs in CD4^+^ Th1 and CD8^+^ Teff subsets.

**(K)** Bubble plot showing enriched antiviral-related signaling pathways in CD4^+^ T cells from VIPA versus Placebo- treated mice.

**(L)** Expression of *Gzma, Ifng,* and *Ccl5* in T cells overlaid on the UMAPs, highlighting enrichment of cytotoxic and effector subsets in VIPA-treated mice.

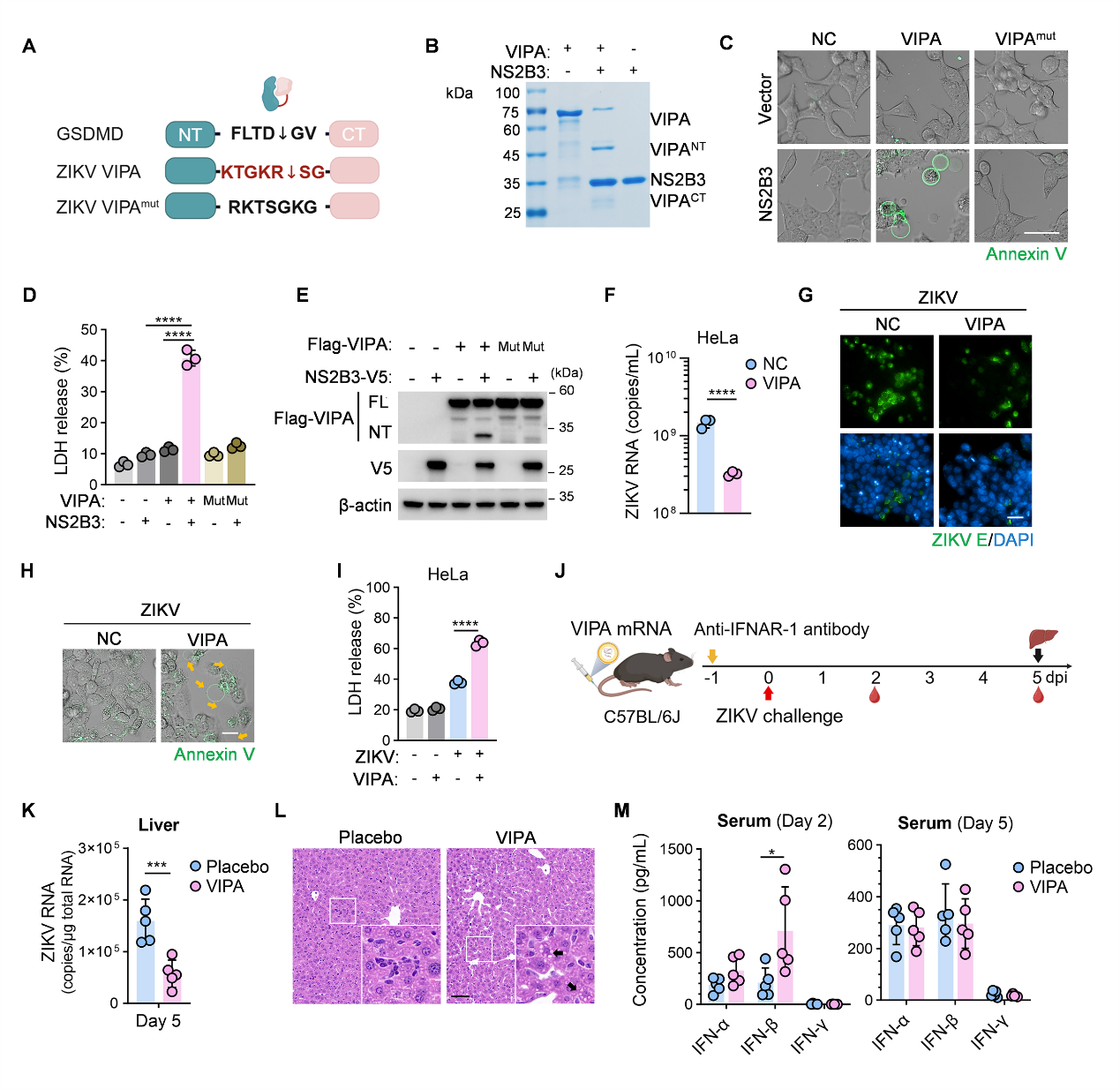

**fig. S4. Design and validation of ZIKV VIPA.**

**(A)** Schematic representation of the ZIKV VIPA and ZIKV VIPA^mut^ constructs, created with BioRender.com.

**(B)** SDS-PAGE showing *in vitro* cleavage of VIPA by ZIKV NS2B3.

**(C to E)** HEK293T cells (n = 3) co-transfected with ZIKV NS2B3 and either VIPA or VIPA^mut^. Live-cell imaging was performed using FITC-Annexin V at 12 h post- transfected **(C)**. LDH release was measured at 12 h post-transfection **(D)**. Western blot analysis of full-length and cleaved VIPA, together with ZIKV NS2B3, was conducted 24 h post-transfection **(E)**. Scale bar, 20 μm.

**(F to I)** HeLa cells (n = 3) were infected with ZIKV and treated with or without VIPA. Quantification of ZIKV RNA copies in HeLa cells infected with ZIKV and treated with or without VIPA at 2 dpi by RT-qPCR **(F)**. Immunofluorescence assays show the level of viral E protein **(G)**. Scale bar, 50 μm. Pyroptotic cell death and cytotoxicity were assessed by live-cell imaging with FITC-Annexin V **(H)**, detection of LDH release at 2 dpi **(I)**. Scale bar, 20 μm.

**(J to M)** C57BL/6 mice (n = 5) treated with VIPA mRNA-LNP following IFNAR-1 blockade and ZIKV infection. **(J)** Experimental schematic created with BioRender.com: yellow arrow indicates antibody-mediated blockade, red arrow indicates viral infection, black arrow indicates experiment termination for tissue analysis. **(K)** Liver viral load measured 5 dpi. **(L)** Representative H&E-stained liver sections at 5 dpi. Scale bar, 25 μm. **(M)** Serum levels of multiple interferons at 2 and 5 dpi.

Data are presented as mean ± SD.

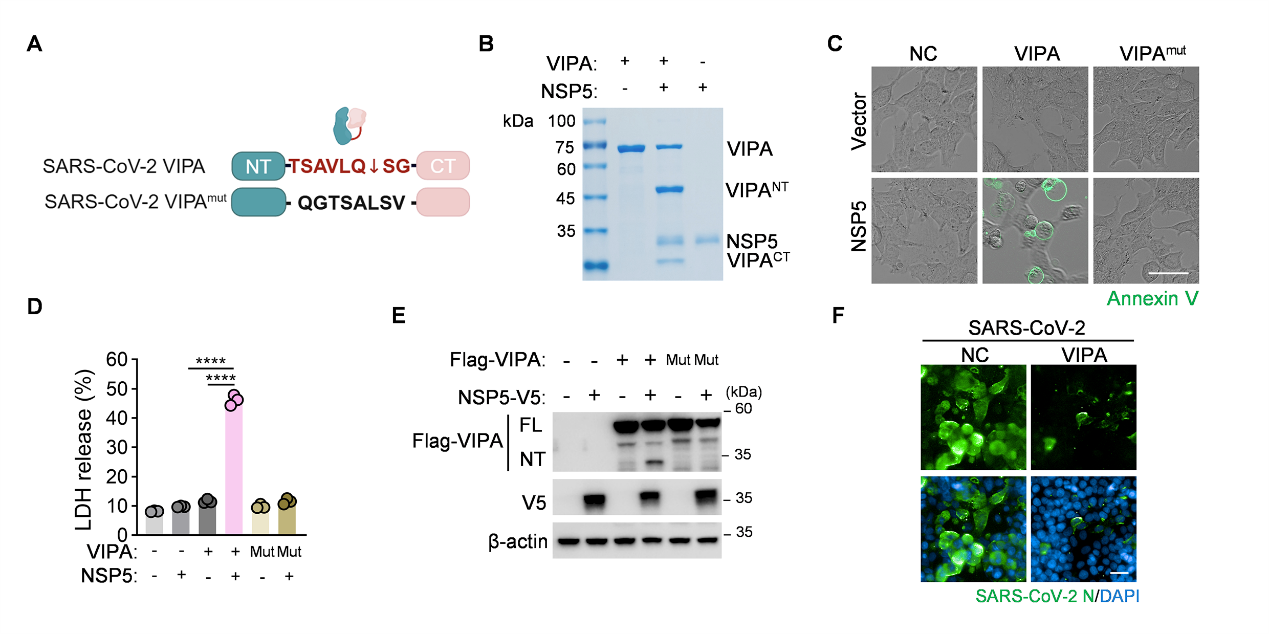

**fig. S5. Design and Validation of SARS-CoV-2 VIPA.**

**(A)** Schematic representation of the SARS-CoV-2 VIPA and SARS-CoV-2 VIPA^mut^ constructs, created with BioRender.com.

**(B)** SDS-PAGE analysis showing in vitro cleavage of VIPA by SARS-CoV-2 NSP5.

**(C to E)** HEK293T cells (n = 3) co-transfected with SARS-CoV-2 NSP5 and either VIPA or VIPA^mut^. **(C)** Live-cell imaging was performed using FITC-Annexin V at 12 h post-transfection (Scale bar, 20 μm). **(D)** LDH release was measured at 12 h post-transfection. **(E)** Western blot analysis of full-length and cleaved VIPA, together with NSP5.

**(F)** HeLa-ACE2 cells (n = 3) transfected with VIPA and infected with SARS-CoV-2. N protein expression in SARS-CoV-2-infected HeLa-ACE2 cells with and without VIPA treatment. Scale bar: 50 μm.

Data are presented as mean ± SD.

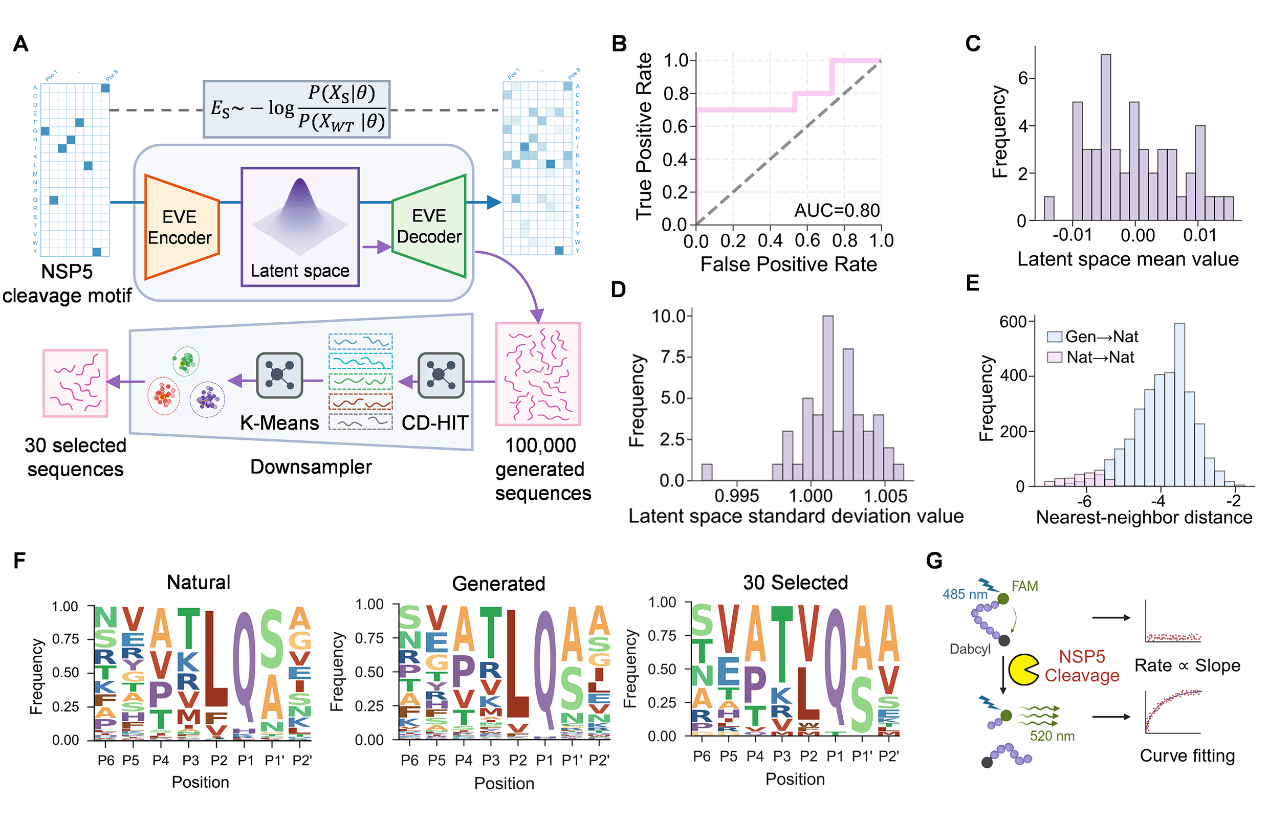

**fig. S6. Evolutionary landscape-guided generation and diversity-aware sampling for VIPA design.**

**(A)** EVE framework and sequence-design workflow. EVE is a variational autoencoder (VAE): one‑hot-encoded input sequences are mapped by the encoder into a latent space, and the decoder outputs softmax probabilities, learning regularities in natural sequences via reconstruction. The evolutionary score (*E_s_*) quantifies evolutionary feasibility as the log‑likelihood ratio of a target sequence relative to the wild type. Design begins by sampling the latent space to decode 100,000 candidates, which are de‑redunded with CD‑HIT and initially filtered by the score; candidates are then embedded in a two‑dimensional sequence space by pairwise similarity and clustered, and centroid‑proximal representatives are selected to yield 30 diverse, high‑activity sequences for experimental evaluation. Created with BioRender.com.

**(B)** ROC curve of the evolutionary score for distinguishing the cell‑death outcome on an independent test set (n = 25).

**(C and D)** Posterior means and standard deviations of training samples in the EVE latent space; means are near 0 and standard deviations near 1, consistent with the model prior.

**(E)** Distributions of generated‑to‑natural (G→N) and natural‑to‑natural (N→N) nearest‑neighbor distances.

**(F)** Sequence logo of the natural, generated, and selected peptides.

**(G)** Schematic of FRET-based in vitro screening for candidate octapeptides to assess NSP5 cleavage efficiency, created with BioRender.com.
